# CRISPR correction of *GBA* mutation in hiPSCs restores normal function to Gaucher macrophages and increases their susceptibility to *Mycobacterium tuberculosis*

**DOI:** 10.1101/2022.11.26.517656

**Authors:** Sivaprakash Ramalingam, Amit Kumar, Stefanie Krug, Harikrishnan Mohan, Desirazu N Rao, William R Bishai, Srinivasan Chandrasegaran

**Author notes:** Contributed equally to this work (Sivaprakash Ramalingam, Amit Kumar).

## Abstract

Gaucher disease (GD) is an autosomal recessive lysosomal storage disorder caused by mutations in the β-glucocerebrosidase (GCase) *GBA* gene, which result in macrophage dysfunction. To investigate whether correction of *GBA* mutations restores normal function to Gaucher macrophages, we performed CRISPR editing of homozygous L444P (1448T→C) *GBA* mutation in Type 2 GD (*GBA*-/-) hiPSCs, which yielded both heterozygous (*GBA*+/-) and homozygous (*GBA*+/+) isogenic lines. Macrophages derived from *GBA*-/-, *GBA*+/- and *GBA*+/+ hiPSCs, were compared for GCase enzymatic activity, motility, and phagocytosis, all of which showed that *GBA* mutation correction restores normal macrophage functions. Furthermore, we investigated whether lysosomal disorders drive susceptibility to *Mycobacterium tuberculosis*, by infecting *GBA*-/-, *GBA*+/- and *GBA*+/+ macrophages with the virulent H37Rv lab strain. The results showed that impaired mobility and phagocytic activity of Gaucher macrophages, correlated with reduced levels of TB engulfment and TB multiplication, supporting the hypothesis that GD may be protective against tuberculosis.

## Introduction

Human β-glucocerebrosidase (GCase) comprises 497 amino acids with four oligosaccharide chains coupled to specific asparagine residues [1]. GCase is a lysosomal hydrolase that breaks down its substrates, glucosylceramide and glucosylsphingosine, into glucose, ceramide, and sphingosine, respectively. Its deficiency causes glucosylceramide and glucosylsphingosine to accumulate within lysosomes of Gaucher macrophages, resulting in lysosomal dysfunction. GD is characterized by lipid-laden Gaucher macrophages that infiltrate bone marrow and other visceral organs [2-4]. Three different clinical forms of GD (Type 1, Type 2 and Type 3) have been identified, of which Type 2 GD is the most severe form. Since GD is a recessive disorder, the mutations occur in both alleles of the *GBA* gene in patients’ cells. While the common N370S mutation is associated with Type 1 GD, the severely destabilizing L444P (1448 T**→**C) mutation is strongly associated with Type 2 and Type 3 GD. Elevated levels of inflammatory mediators (TNFα, IL-6 and IL-1β) have been reported in the serum of GD patients [3, 4], and GD macrophages have been shown to have migratory defects [5, 6]. Type 1 GD, which occurs at high frequency in Ashkenazi Jews (carrier rate ∼1 in 15), is thought to have originated 800-1400 years ago [7, 8], and the persistence of this and other types of GD mutations in human populations at relatively high levels has prompted the concept that GD mutations may confer a selective advantage. One theory holds that GD homozygosity and/or heterozygosity may be protective against common lethal human infections such as tuberculosis [9, 10].

## Results and Discussion

We had previously reported TALEN-mediated generation of Type 2 homozygous L444P (1448 T**→**C) GD hiPSCs [11]. CRISPR/Cas9 editing of L444P mutations in GD (*GBA*-/-) hiPSCs using sgRNA and a 100 bp single-strand oligonucleotide as the donor template, yielded isogenic lines with heterozygous (*GBA*+/-) and homozygous (*GBA*+/+) gene correction (**Fig. 1a**). Since L444P mutation results in a *Nci* I restriction site, we screened for *GBA*+/- and *GBA*+/+ single cell colonies by digesting ∼800 bp PCR-amplified fragment surrounding the mutant locus with *Nci* I (**Fig. 1b**). Sequencing of the PCR-amplified DNA further confirmed the genotypes of *GBA*-/-, *GBA*+/- and *GBA*+/+ hiPSCs (**Fig. 1c**). Expression of pluripotency markers in *GBA*+/- and *GBA*+/+ hiPSCs, was confirmed by immunostaining using antibodies for *Oct4, Sox2, Tra-1-60, Tra-1-81* and DAPI, respectively. Karyotyping of the *GBA*+/- and *GBA*+/+ hiPSCs, established that the cells were all normal (46XY) without any chromosomal abnormalities, like the GD hiPSCs that were characterized previously [11]. Sequencing of eight sites closely related to the sgRNA target (SI: xls) in *GBA*+/- and *GBA*+/+ hiPSCs, did not reveal any indels that were induced by NHEJ, ruling out off-target cleavage by CRISPR/Cas9 (data not shown).

**Figure 1.**
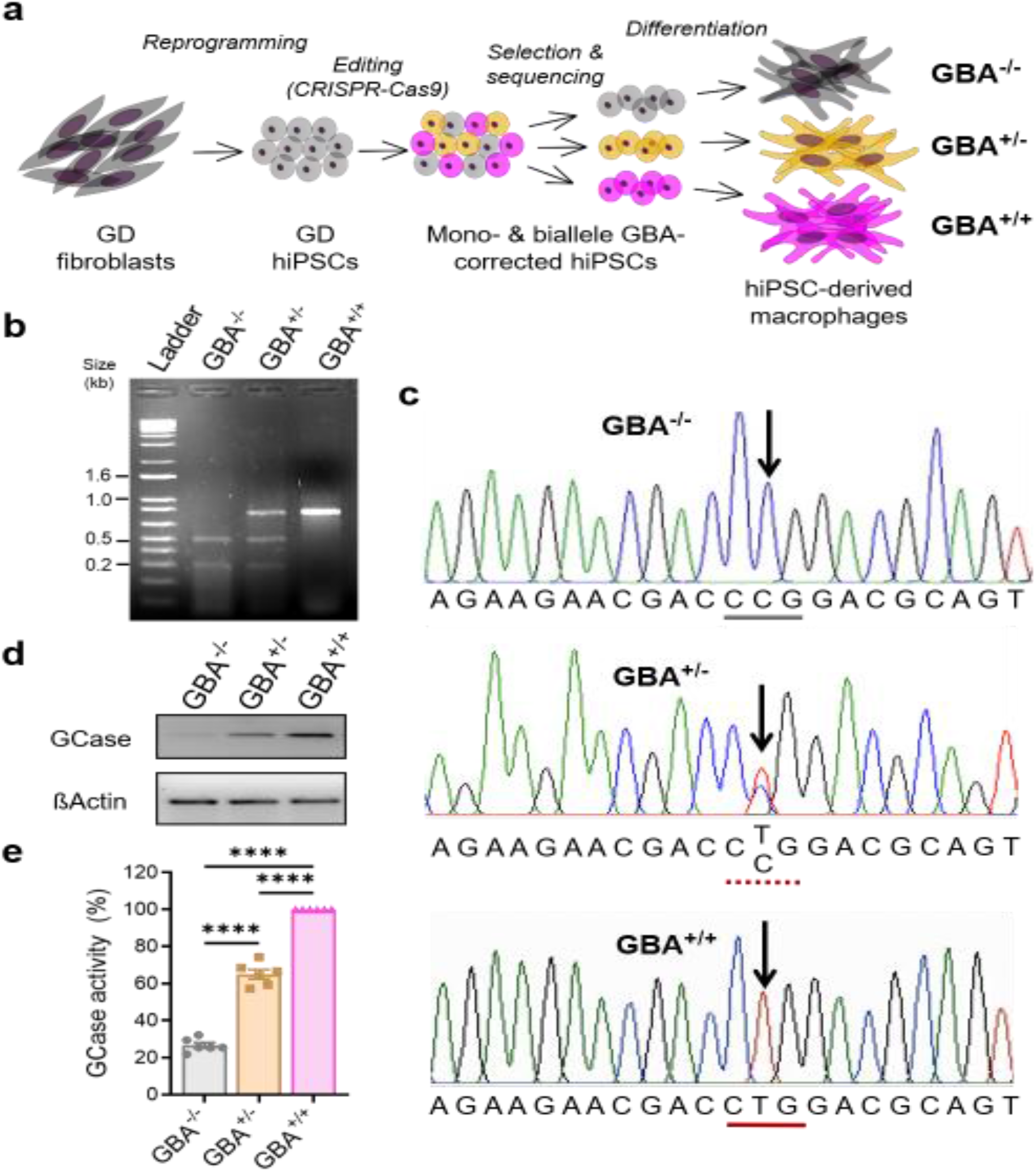
CRISPR correction of *GBA* mutations in GD hiPSCs restores normal GCase activity to Gaucher macrophages. a) Schematic diagram showing generation and genetic correction of GD hiPSCs and their differentiation into macrophages. b) Genotype characterization of GBA-/-, GBA+/- and GBA+/+ hiPSCs by *Nci* I restriction enzyme digest. c) Sequence profiles of the *GBA* mutation locus in *GBA*-/-, *GBA*+/- and *GBA*+/+ hiPSCs. d) Western blot analysis of GCase protein levels. e) GCase enzymatic activity relative to GBA+/+ cells using 4-methylumbelliferyl β-D-glucopyranoside as the substrate: (GBA+/+) > (GBA+/-) > (GBA-/-). ****, p<0.0001.

We differentiated the *GBA*-/-, *GBA*+/- and *GBA*+/+ isogenic hiPSCs, first into monocytes, and then into macrophages using standard protocols (**Fig. 1a**). Western blot analysis using a GCase antibody, confirmed that GCase expression is partially restored in *GBA*+/-, and fully restored in *GBA*+/+ macrophages (**Fig. 1d**). We also monitored the GCase activity directly in *GBA*-/-, *GBA*+/- and *GBA*+/+ macrophages, using 4-methylumbelliferyl-β-D-glucosylpyranoside as the substrate (**Fig. 1e**). The results confirmed partial restoration of GCase activity in *GBA*+/-, and full restoration in *GBA*+/+ compared to *GBA*-/- macrophages.

To determine whether Gaucher macrophages show abnormal chemokine activation and whether their functional defects could be reversed by L444P mutation correction, we examined the expression profiles of TNFα, IL-6, IL-1β and IL-10 in isogenic *GBA*-/- *GBA*+/- and *GBA*+/+ macrophages (**Fig. 2a**). Significant higher levels of IL-1β expression were observed in homozygous *GBA*-/- and heterozygous *GBA*+/- macrophages compared to *GBA*+/+ GBA- corrected macrophages. IL-1β is an important inflammatory mediator that is involved in cell proliferation, differentiation, and apoptosis. Increased risk of cancers, autoimmune disease, and infections associated with human GD, could be attributed to the elevated production of IL-1β by Gaucher macrophages. Like the IL-1β, expression of the proinflammatory cytokines TNFα and IL-6, expression routinely trends higher in isogenic Gaucher macrophages while anti-inflammatory IL-10 appears to be unaffected by *GBA* mutations. Our observations are consistent with previous studies reporting elevated TNFα, IL-6, and IL-1β expression in Gaucher macrophages differentiated from patient-derived hiPSCs with Type 1, 2, and 3 GD, compared to non-isogenic control cells [2, 3].

**Figure 2.**
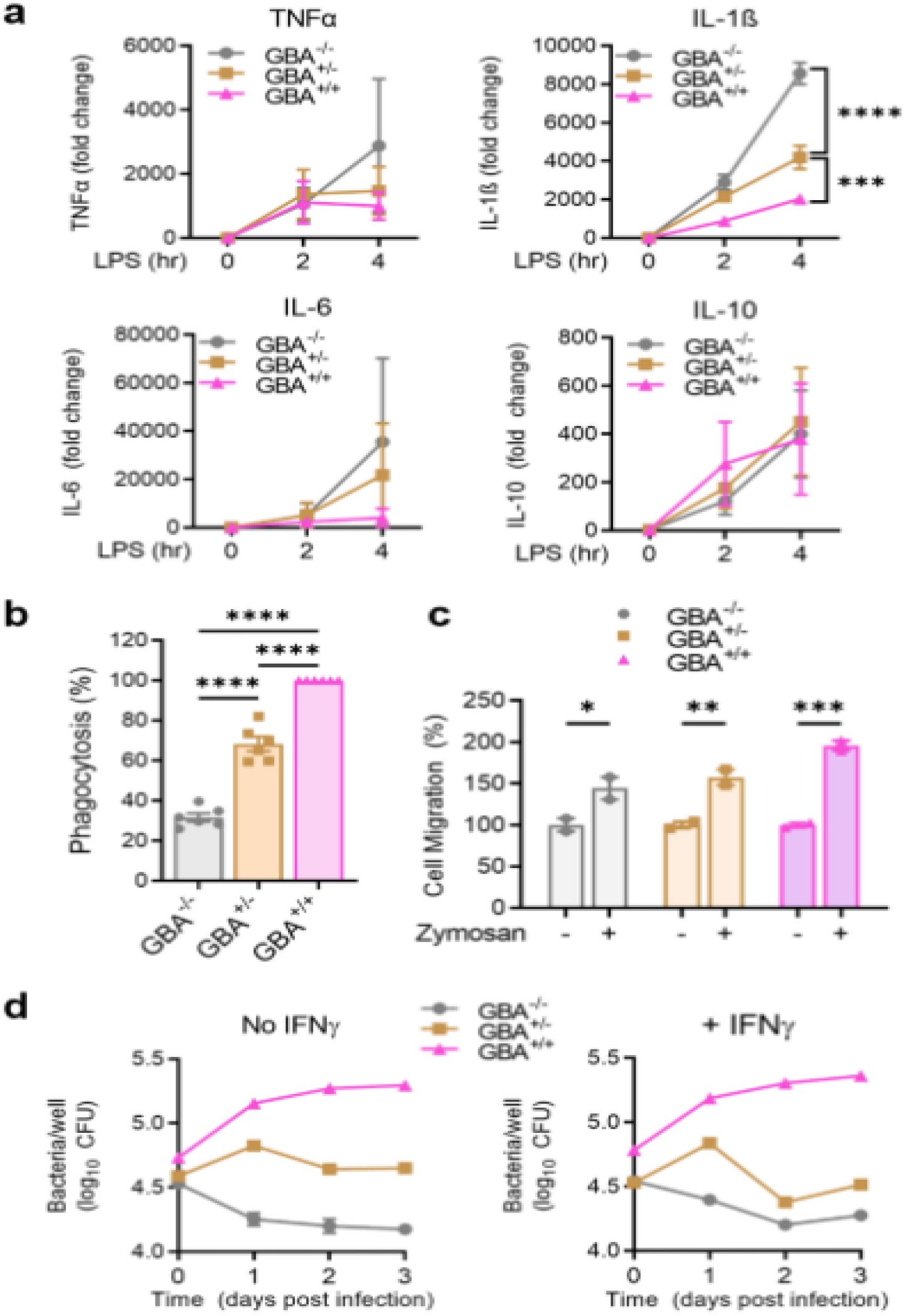
CRISPR-Cas9 correction of GBA mutation in GD hiPSCs restores normal phagocytic, motility and immune functions and increases susceptibility of Gaucher macrophages to *M. tuberculosis*. a) TNFα, IL-6, IL-1β and IL-10 transcript levels in isogenic GBA-/-, GBA+/- and GBA+/+ macrophages stimulated with LPS for up to 4h. Significantly higher levels of IL-1β expression were observed in Gaucher macrophages: (GBA-/-) > (GBA+/-) > (GB+/+). b) Phagocytic activity of macrophages using zymosan particles relative to phagocytosis in GBA+/+ macrophages. c) Analysis of macrophages cell motility by transwell migration assay. (d) Analysis of macrophage susceptibility to *M. tuberculosis* H37Rv infection and intracellular replication. Graphs show intracellular bacterial burden after infection in resting (left, “No IFNy”) or primed (right, “+ IFNy”) macrophages. *, p<0.05. **, p<0.01. ***, p<0.001. ****, p<0.0001.

Next, we examined whether CRISPR editing of L444P mutations in GD hiPSCs, restores normal phagocytic function and motility to Gaucher macrophages. We evaluated phagocytic functionality of GD (*GBA*-/-), *GBA*+/- and *GBA*+/+ macrophages by zymosan particle engulfment assay (**Fig. 2b**). As expected, GD macrophages showed only minimal engulfment of zymosan particles while phagocytosis was moderately recovered in *GBA*+/- macrophages. In contrast, biallelic *GBA*-corrected macrophages exhibited maximal engulfment of zymosan particles, indicating L444P mutation correction restores normal phagocytic potential to Gaucher macrophages.

Since migration is critical for macrophage immune function, we compared the mobility of *GBA*-/-, *GBA*+/- and *GBA*+/+ macrophages by transwell migration assay (**Fig. 2c**). Like phagocytosis phenotypes, we observed that the migratory defect of *GBA*-/- macrophages was partially restored in *GBA*+/- macrophages, while biallelic *GBA* correction (*GBA*+/+) fully restored cell motility compared to GD macrophages. Promisingly, our findings suggest monoallelic L444P mutation correction may sufficiently enhance cell migration of Gaucher macrophages for a functional immune response, which is consistent with GD being a recessive lysosomal storage disorder.

The isogenic hiPSC-derived GD and *GBA*-corrected macrophages offer an excellent model to investigate whether lysosomal disorders drive susceptibility to *M. tuberculosis* and, whether *GBA* mutation correction restores normal lysosomal functions, *M. tuberculosis* susceptibility and infectivity to Gaucher macrophages. *M. tuberculosis* replicates in macrophages by inhibiting phagosome-lysosome fusion. Lysosomal dysfunction might prevent the formation of stable TB granulomas. leading to secondary necrosis and altered tuberculosis susceptibility in GD patients. To determine how GD and *GBA* mutation correction affect infectivity and growth of *M. tuberculosis* in macrophages, we infected the isogenic *GBA*-/-, *GBA*+/- and *GBA*+/+ macrophages with *M. tuberculosis* H37Rv under BSL3 conditions and monitored the intracellular bacterial burden within these cells (**Fig. 2d**). Surprisingly, the cellular environment of GD macrophages appears to impair the uptake and intracellular replication of H37Rv. In contrast, biallelic *GBA*-corrected macrophages supported robust H37Rv infection and growth while bacterial replication was static in *GBA*+/- macrophages. Similar results were obtained for both untreated and IFNγ-activated macrophages (**Fig. 2d**). Together, our findings suggest GD may confer a level of protection against tuberculosis and that *GBA* mutation correction increases Gaucher macrophage susceptibility to *M. tuberculosis*.

These findings lend credibility to the hypothesis that *GBA*-/- homozygosity and *GBA*+/- heterozygosity are protective against tuberculosis, and this may account in part for the persistence of mutations in human populations. Recently, Fan et al. [12] showed that GD-/- zebrafish are resistant to infection with either *M. tuberculosis* or *M. marinum* (a closely related mycobacterial species that is a natural fish pathogen) [13]. They further showed that this resistance is mediated by direct anti-bacterial activity by lysosomal glucosylsphingosine, which accumulates in *GBA*-/- macrophages to high levels. Interestingly, in contrast to our findings in which *GBA*+/- human macrophages showed intermediate resistance to *M. tuberculosis* infection, *GBA*+/- heterozygous zebrafish in their study showed equivalent susceptibility to mycobacterial infection to that seen in wild-type fish. One possible explanation for this disparate observation is that Fan et al. [12] studied the N370S mutation in zebrafish, commonly observed in Ashkenazi Jews, which causes mild disease (non-neuropathic, Type 1) and modest life expectancy defects in homozygotes and is associated with reduced but not abolished GCase enzymatic activity. In contrast, our work was done with hiPSCs carrying L444P mutation in heterozygotes, which drastically reduces GCase activity and causes severe disease (neuropathic, Type 2) in homozygotes that leads to mortality in early childhood.

In summary, we demonstrate that targeted CRISPR correction of a severe *GBA* mutation in hiPSCs, restores normal functions to Gaucher macrophages. In addition, our findings support the hypothesis that GD confers protection against tuberculosis, consistent with a recent report using the zebrafish model [12]. Promisingly, our study suggests that it might be feasible to develop either hematopoietic (HSC) or CD34+ stem cells-based gene therapy as a permanent curative alternative to the expensive life-long GCase enzyme replacement therapy, to treat Type 1 GD [14, 15].

## Materials and Methods

Experimental protocols are described in detail in Supplementary Information: Extended Materials and Methods.

## Acknowledgments

This work was supported by a grant (AI133530) from NIH. Dr. Amit Kumar was partially supported by a Research Demonstration Project grant from the CF Foundation.

## Author Contributions

S.C. and W. B. conceptualized the study. S.C., S.R., W.B. and A.K. designed the experiments. S.R. performed functional characterization GD hiPSCs. H.M. performed Cas9 off-target cleavage analyses. A.K. performed macrophage phagocytosis, motility, and *M. tb* infectivity assays. S.K. and A.K. performed data analyses and prepared the figures. S.C. and W.B. wrote the manuscript. D.R. provided advice and critical reading of the manuscript. S.C. and W.B. provided lab supervision. All authors reviewed and edited the manuscript.

